# The Evolution of Dynamic and Flexible Courtship Displays That Reveal Individual Quality

**DOI:** 10.1101/2022.09.13.507802

**Authors:** Sam H. Hollon, Irene García Ruiz, Thor Veen, Tim W. Fawcett

## Abstract

Sexual selection is a major force shaping morphological and behavioral diversity. Existing theory focuses on courtship display traits such as morphological ornaments whose costs and benefits are assumed be to fixed across individuals’ lifetimes. In contrast, empirically observed displays are often inherently dynamic, as vividly illustrated by the acrobatic dances, loud vocalizations, and vigorous motor displays involved in courtship behavior across a broad range of taxa. One empirically observed form of display flexibility occurs when signalers adjust their courtship investment based on the number of rival signalers. The predictions of established sexual selection theory cannot readily be extended to such displays because display expression varies between courtship events, such that any given display may not reliably reflect signaler quality. Thus, we lack an understanding of how dynamic displays coevolve with sexual preferences and how signalers should tactically adjust their display investment across multiple courtship opportunities. To address these questions, we extended an established model of the coevolution of a female sexual preference and a male display trait to allow for dynamic, flexible displays. We find that a dynamic display can coevolve with a sexual preference away from their naturally selected optima, though display intensity is a weaker signal of male quality than for static ornaments. Furthermore, we find that males evolve to decrease their display investment when displaying alongside more rivals. This research represents a first step towards generalizing the findings of sexual selection theory to account for the ubiquitous dynamism of animal courtship.

**Significance Statement:** Animal courtship displays are typically costly for survival: songs attract predators; dances are exhausting; extravagant plumage is cumbersome. Because of the trade-off between mating benefits and survival costs, displaying individuals often vary their displays across time, courting more intensely when the potential benefit is higher or the cost is lower. Despite the ubiquity of such adjustment in nature, existing theory cannot account for how this flexibility might affect the coevolution of displays with sexual preferences, nor for the patterns of tactical display adjustment that might result, because those models treat displays as static, with fixed costs and benefits. Generalizing a well-studied model of sexual selection, we find that a static display and a flexible display can evolve under similar conditions. Our model predicts that courtship should be less intense when more competitors are present.

## 1 Introduction

Sexual selection is a major force shaping morphological and behavioral diversity (Andersson, 1994). A vast body of theoretical work has uncovered conditions under which sexual preferences can lead to the exaggeration of conspicuous morphological “ornaments” that impose survival costs on the individuals that bear them (Kuijper et al., 2012). Established models assume these ornaments to be static such that an individual’s display expression changes only on long timescales (e.g., across life stages or through growth), if at all. Yet empirically observed displays are often inherently dynamic, as vividly illustrated by the acrobatic dances, loud vocalizations, and vigorous motor displays involved in courtship behavior across a broad range of taxa (Byers et al., 2010).

In contrast to the ornaments of established sexual selection theory, these dynamic displays vary in their expression both within and between courtship events (Patricelli et al., 2016). The ubiquity of such displays poses a major theoretical challenge because established models of sexual selection rely on a fixed relationship between an individual’s quality and display expression: for every possible individual quality, there exists a unique display value, from which we may determine an associated mating benefit and survival cost. Then a preference for a costly display trait may evolve if the display intensity is an honest signal of quality such that individuals of higher quality have consistently higher-intensity displays than those of lower quality. This honesty may be maintained by a handicap mechanism that ensures that the fitness of higher-quality individuals increases more with additional investment in the display (Getty, 2006; van Doorn & Weissing, 2006). In contrast, when display intensity is dynamic or flexible, any given display may not reflect signaler quality. In particular, with a flexible display, variation in display intensity could conceivably undermine signal honesty if individuals of lower quality secure a greater mating advantage than those of higher quality by investing heavily in the display under some conditions and little under others.

No existing model of which we are aware addresses this possibility. Hutchinson et al. (1993) constructed a model of song displays in which a female preference led males to vary the timing of their singing throughout the day. However, the model assumed a fixed female preference for more intense displays. Thus, the model does not shed light on how a preference for a dynamic display trait might arise nor on the patterns of signaling that would be produced as the preference and display coevolve. Similarly, South et al. (2012) showed that a male bias towards courting higher-fecundity females can evolve when females prefer males who invest more in courtship, demonstrating another route to the evolution of signal flexibility. But once again, the female preference was assumed by the modelers, not free to evolve, eliminating the coevolutionary dynamics of present interest.

To fill this theoretical gap, we investigate two intertwined questions: (1) Can a dynamic, flexible display coevolve with a sexual preference for that display through a handicap mechanism? (2) If so, how should individuals tactically adjust their investment in the display in response to competition from rival signalers?

We begin by giving sharp definitions to *static, dynamic*, and *flexible* displays. We then construct an individual-based simulation of the coevolution of a male static display trait with a female sexual preference, following the standard assumptions of sexual selection theory. Specifically, this model relies on a revealing-handicap mechanism (Maynard Smith, 1985), which is well-studied in the case of static displays (van Doorn & Weissing, 2006). Under this handicap, for the same investment in the display, males of higher and lower quality pay the same costs, but the former are more attractive to females that have a preference for the display.

We then extend this established model by implementing a display that is dynamic and flexible, rather than static, and a corresponding preference. The display is dynamic in that males express it (and pay its associated cost) only when they actively court a female. The display is flexible in that males may adjust their display investment in response to rival males that they display alongside (who are competing to mate with the same female). There are two reasons that flexibility based on the number of rivals is a compelling avenue for research. First, there is existing empirical evidence of this form of flexibility, for example in fiddler crabs, *Austruca mjoebergi*, which wave their major claw faster as the level of competition increases (Milner et al., 2012). Second, it is not obvious through intuition alone whether males should, in general, display more or less when they display alongside more rivals. When rival males engage in costly fighting and those costs are sufficiently high, we would expect males to signal less when they face formidable rivals. For example, lekking sage grouse, *Centrocercus urophasianus*, delay signaling to avoid aggression from rivals (Patricelli et al., 2016). But what about when competition between males acts only through female choice, not direct contests? On the one hand, when a male faces more rivals, he might have to signal more intensely to capture a female’s attention and so obtain a reasonable chance of being chosen to mate. On the other hand, if competition is too intense, it might be better to reduce display effort, or stop displaying altogether. It is precisely this difficulty of giving a satisfying narrative account of the evolutionary outcome that necessitates formal modeling (Otto & Rosales, 2020).

## 2 Static, Dynamic, and Flexible Displays

We distinguish displays along two lines: static *versus* dynamic, and non-flexible *versus* flexible.

We define *static displays* as display traits whose intensity is constant on long timescales, normally throughout an individual’s adult lifetime or throughout stages of growth (Figure 1a). An important property of purely static displays is that they do not require us to distinguish between when an individual is actively displaying and when it is being passively observed, such that the associated costs are independent of how often it encounters potential mates. We can then reasonably model a display as a constant trait value determined by the individual’s quality in advance of any encounters with potential mates. Biologically, a purely static display can be thought of as a morphological trait that is displayed similarly at all times. For example, a bird’s plumage would be purely static if the individual never adjusted the plumage’s appearance, such as through a dance or strutting, to catch the eye of a receiver.

**Fig. 1:**
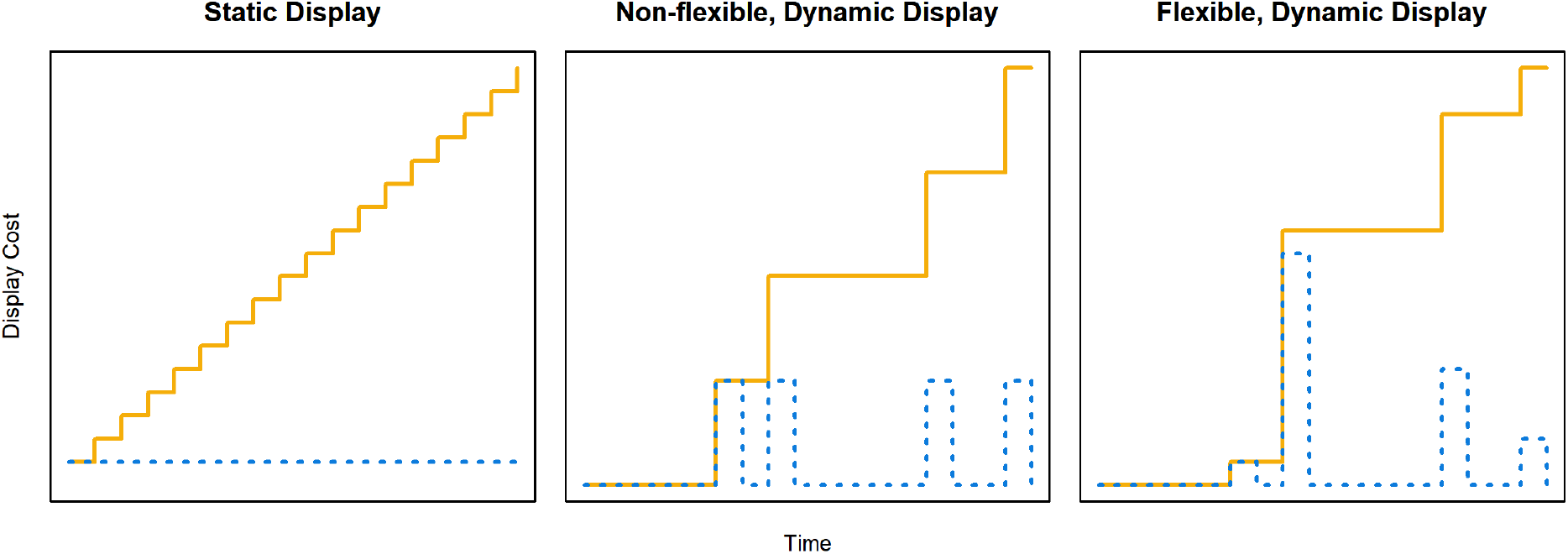
The costs of three types of display. The blue dotted lines show the survival cost of a hypothetical display at each time step during an individual’s adult lifetime for each type of display, while the yellow lines show the cumulative lifetime cost. The static display’s intensity is constant across time, so the cumulative cost (and benefit, not shown) increases linearly (left). Both the non-flexible, dynamic (center) and flexible, dynamic (right) displays’ intensities, and hence the costs (and benefits, not shown), are zero except at four time steps when the individual actively displays. However, for the non-flexible display, the cost is the same for each display event, whereas for the flexible display, the cost for each display event can differ, due to varying display intensity across display contexts. Note that until the first display event, the cost of the dynamic displays is zero.

We define *dynamic displays* as display traits whose intensity is negligible except when the signaler actively displays. In contrast to that of a static display, the cost and benefit of a dynamic display depends on the display’s use; with a purely dynamic display, a signaler that never has the opportunity to display pays no survival cost and gains no mating benefit, regardless of what its investment in the display would have been. For example, birdsong is dynamic in that the bird pays the cost of displaying, such as increasing attention from predators, only while singing and can stop singing to cease paying that cost while also ceasing to receive the benefit.

Following Wainwright et al. (2008), we define *flexible displays* as display traits whose intensity, and consequently their cost and benefit, varies across different signaling contexts. Such variation is common empirically. For example, male house flies, *Musca domestica*, adjust their displays according to the differing preferences of potential mates (Meffert & Regan, 2002); male Australian terrestrial toadlets, *Pseudophryne bibronii*, increase their call rate in the presence of a female odor, when the cost-effectiveness of displaying is higher (Byrne & Keogh, 2007); male black widows, *Latrodectus hesperus*, selectively display when the risk of sexual cannibalism is lower (Baruffaldi & Andrade, 2015); and male fiddler crabs adjust the rate they wave their claws for females based on the number of rival males waving concurrently (Milner et al., 2012).

Flexible displays are a subset of dynamic displays: because context can alter signal intensity on short timescales, flexible displays are inherently dynamic. As such, there are three possible types of display in our typology: (1) static displays; (2) non-flexible, dynamic displays; and (3) flexible, dynamic displays (Figure 1). However, few real-world displays are likely to be well approximated as purely static or dynamic or as purely flexible or non-flexible (Patricelli et al., 2016); this is merely a simplified representation to make models more tractable. In reality, displays may consist of multiple components, each with varying degrees of dynamism depending on the timescale over which they are adjusted, from very rapid movements during a single courtship event to seasonal or even longer-term changes. The components may also interact with each other: for example, shiny plumage may be visible at all times, but during active courtship it becomes more salient both to potential mates and to predators. Thus, explaining most of the diversity of animal courtship displays requires accounting for dynamism and flexibility.

## 3 Model

We model a population of *N* individuals with non-overlapping, discrete generations and an even primary sex ratio. Each individual has autosomal, diploid genetic values for (1) quality, (2) investment in a static display, (3) investment in a dynamic display, (4) sexual preference for the static display, and (5) sexual preference for the dynamic display. These traits are summarized in Table 1. Quality is expressed in all individuals, while display expression is limited to males and preference expression is limited to females.

**Table 1:** Parameter values and genetic values for all individual-based simulations. An individual’s genotype is completely described by these trait values. The dynamic display function *t_d_*(*n*) is not listed, as it is a phenotypic value only, completely determined by the genetic values *d*_1_ and *d*_2_. Likewise, genetic viability *v* is not shown because it is determined by quality *q*.

| Parameter | Value | Meaning |
| --- | --- | --- |
| $N$ | 1000 | Population size |
| $\bar{n}$ | 3 | Mean number of rival males |
| $\alpha$ | 0.001 | Cost of preferences |
| $\beta_s$ | 0.15, 0.25 | Cost of static display |
| $\beta_d$ | 0.015, 0.025 | Cost of dynamic display |
| $\gamma$ | 0.6 | Probability that mutations in genetic viability are downward |
| $\mu$ | 0.05 | Mutation rate per trait per offspring |
| $\sigma$ | 0.001 | Mutation standard deviation |
| Genetic Trait |  | Meaning |
| $d_1$ | | Dynamic display intercept |
| $d_2$ | | Dynamic display slope |
| $p_s$ | | Preference for static display |
| $p_d$ | | Preference for dynamic display |
| $q$ | | Quality |
| $t_s$ | | Investment in static display |

### 3.1 Traits

#### 3.1.1 Quality

Quality *q* incorporates all genetic factors other than display and preference expression that affect viability. *q* may assume any real value, with *q* = 0 corresponding to optimum genetic viability. We assume that genetic viability v declines exponentially with *q* according to *v* = exp(−|*q*|).

#### 3.1.2 Static Display

Males make a one-time investment in their static display during development that reduces their survival to maturity and determines their static display investment *t_s_* as an adult. For example, *t_s_* could represent the size of a morphological ornament that is fixed at the end of development. For ease of biological interpretation, *t_s_* is restricted to non-negative values, with a value of 0 corresponding to no ornament and larger values corresponding to larger ornaments.

#### 3.1.3 Dynamic Display

The dynamic display function *t_d_*(*n*) gives a male’s investment in the dynamic display when displaying alongside *n* rival males. Because there is little empirical evidence on the functional form that *t_d_* might take in nature, we assume for simplicity that *t_d_* is linear in *n*. We constrain the display investment to non-negative values, yielding

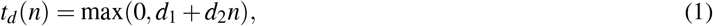

where the intercept *d*_1_ corresponds to the non-flexible part of the investment in the display and the slope *d*_2_ corresponds to the flexible part. Intuitively, a male invests a baseline effort of *d*_1_ when displaying in the absence of rivals and adjusts this investment upwards, downwards, or not at all according to *n* and the sign of *d*_2_. If *d*_2_ > 0, the male displays more alongside more rivals; if *d*_2_
< 0, he displays less. When *d*_2_ = 0, the display is dynamic (it is only expressed during courtship events) but non-flexible (it is always expressed at the same intensity), whereas when *d*_2_ ≠ 0, the display is both dynamic and flexible (its intensity varies between display events).

#### 3.1.4 Preferences

For simplicity, we assume that female preferences are fixed across their lifetime. Let *p_s_* be a female’s preference for the static display and *p_d_* her preference for the dynamic display. A positive value of *p_s_* or *p_d_* implies a preference for males with higher display intensity, while a negative value implies a preference for males with lower display intensity. A value of zero implies no preference, i.e. random mating.

### 3.2 Procedure

Each generation, we simulate three processes in sequence: mortality, nonrandom mating, and reproduction. Figure 2 illustrates the simulation procedure for each generation.

**Fig. 2:**
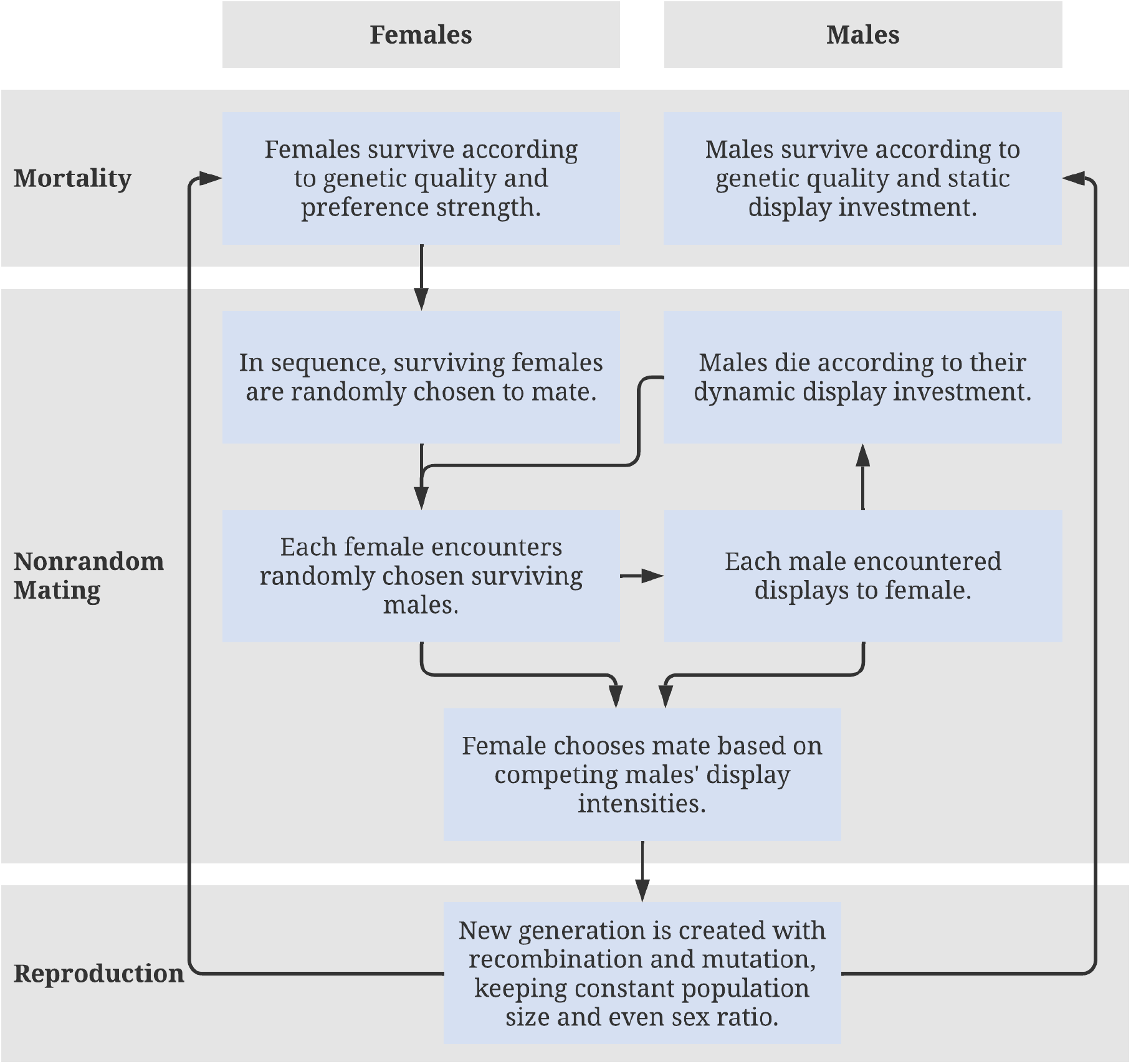
Diagram of the simulation procedure executed each generation.

#### 3.2.1 Mortality

At the start of the generation, each individual survives according to its initial viability, as determined by its genotype. Female viability *S_f_* increases with genetic viability *v* and decreases with preferences *p_s_* and *p_d_* according to

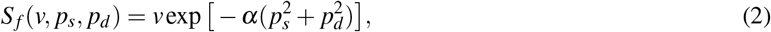

where the cost parameter *α* determines how quickly viability decreases with increasing preference. Similarly, male viability *S_m_* increases with genetic viability *v* and decreases with static display expression *t_s_* according to

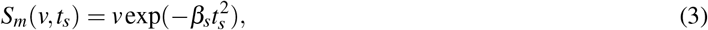

where the cost parameter *β_s_* determines how quickly a male’s viability decreases with investment in the static display. This expression for male viability accounts for survival to adulthood, before any courtship. In contrast to the static display cost, the dynamic display cost has no effect on this initial viability. Instead, the dynamic display cost reduces viability after each display, as described below.

#### 3.2.2 Nonrandom Mating

Next, we randomly draw one female *N* times, with replacement, from the surviving population. In sequence, each female drawn randomly samples *n* + 1 surviving males, where *n* is a Poisson-distributed random number with a mean of 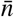. (Note that we add the 1 to this number to ignore cases where a female samples zero males) Thus, the mean number of *rival males* sampled, that is, the number of males in addition to any focal male in a sample, is 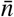.

Each of the sampled males then displays to the female, with an investment of *t_s_* in the static display and *t_d_*(*n*) in the dynamic display. We constrain the investment in each display to non-negative values for ease of interpretation.

A male’s attractiveness *A*, as perceived by a female with preferences *p_s_* and *p_d_*, is given by

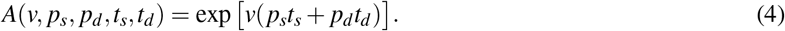

Note that attractiveness increases more with display investment for males of higher viability. The model thus implements a revealing handicap, following the implementation of van Doorn and Weissing (2006). The biological interpretation is that, for the same level of investment in the display, the realized display intensity is greater for males of higher quality. For example, if a male is otherwise more vigorous, then the same investment in a motor display might correspond to a more skillful motor display. The choosing female then selects and mates with one of the displaying males with a probability proportional to *A*.

After displaying and possibly being selected by the female to mate, each male survives with probability 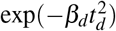, where the cost parameter *β_d_* determines how quickly survival decreases with dynamic display investment in the current courtship attempt. Biologically, this survival risk could represent the display attracting the attention of predators. Thus, by investing in the dynamic display, a male increases his probability of being chosen by the current female (assuming *p_d_ >* 0 for that female) but reduces his probability of surviving to be sampled by future females.

#### 3.2.3 Reproduction

Each mating produces one offspring. We determine the genetic values of the offspring through standard Mendelian inheritance, assuming no genetic dominance and that all loci are unlinked.

Following established models of sexually selected handicaps (Iwasa et al., 1991; van Doorn & Weissing, 2006), we assume that each trait mutates with probability μ in each offspring and that mutations in displays and preferences are unbiased, while mutations in quality are downward-biased (i.e., *q* is more likely to decrease than increase). If a mutation occurs in a display or preference trait, we simulate the mutation by adding a normal deviate with a mean of zero and a small standard deviation σ to the genetic value. If a mutation occurs in quality, we first compute the magnitude of the mutation as a normal deviate of mean zero with standard deviation σ, then we determine the sign of the change: the mutation is downward (negative) with probability *γ*, where *γ* > 0.5, and upward with probability 1 – *γ*.

### 3.3 Simulations

To separate the effects of dynamism and flexibility in display expression, we run three sets of simulations. First, we analyze the results of the baseline static-display model. Second, we replace the static display with a dynamic display but require that males invest the same effort in every display they give (i.e., the display is dynamic but non-flexible). Third, we incorporate flexibility by allowing males to vary their investment in the dynamic display in response to the number of rivals they face. To reveal the effects of selection more clearly, in all simulations, we assume a preexisting female preference for the display, initially holding the preference constant at 0.3 while allowing the male display to evolve from 0. Then, once evolutionary change in the display investment has slowed down, we allow the female preference to coevolve with the male display. Similarly, to isolate the effects of display flexibility in the simulations that include the dynamic, flexible display, we initially hold the display slope *d*_2_ constant, allowing only the display intercept *d*_1_ to evolve from its initial value, before allowing the two display components to coevolve with one another and the female preference.

## 4 Results

### 4.1 Static Display

The first set of simulations included only the static display, with the dynamic display fixed at *t_d_* = 0. For these simulations, we set the initial values for the preference and display to *p_s_* = 0.3 and *t_s_* = 0, respectively, for all individuals in the first parental generation. For these runs, we set the dynamic display cost *β_s_* to two values, 0.15 and 0.25, and ran 50 replicates for each value. We set the values of all other parameters as in Table 1.

The static display and preference coevolved to positive stable values (Figure 3a), in line with the results of established sexual selection models (Kuijper et al., 2012). Also as expected, once the two traits neared these values, we observed a positive relationship between the two traits: the display evolved to higher values when the preference was high (Figure 3b).

**Fig. 3:**
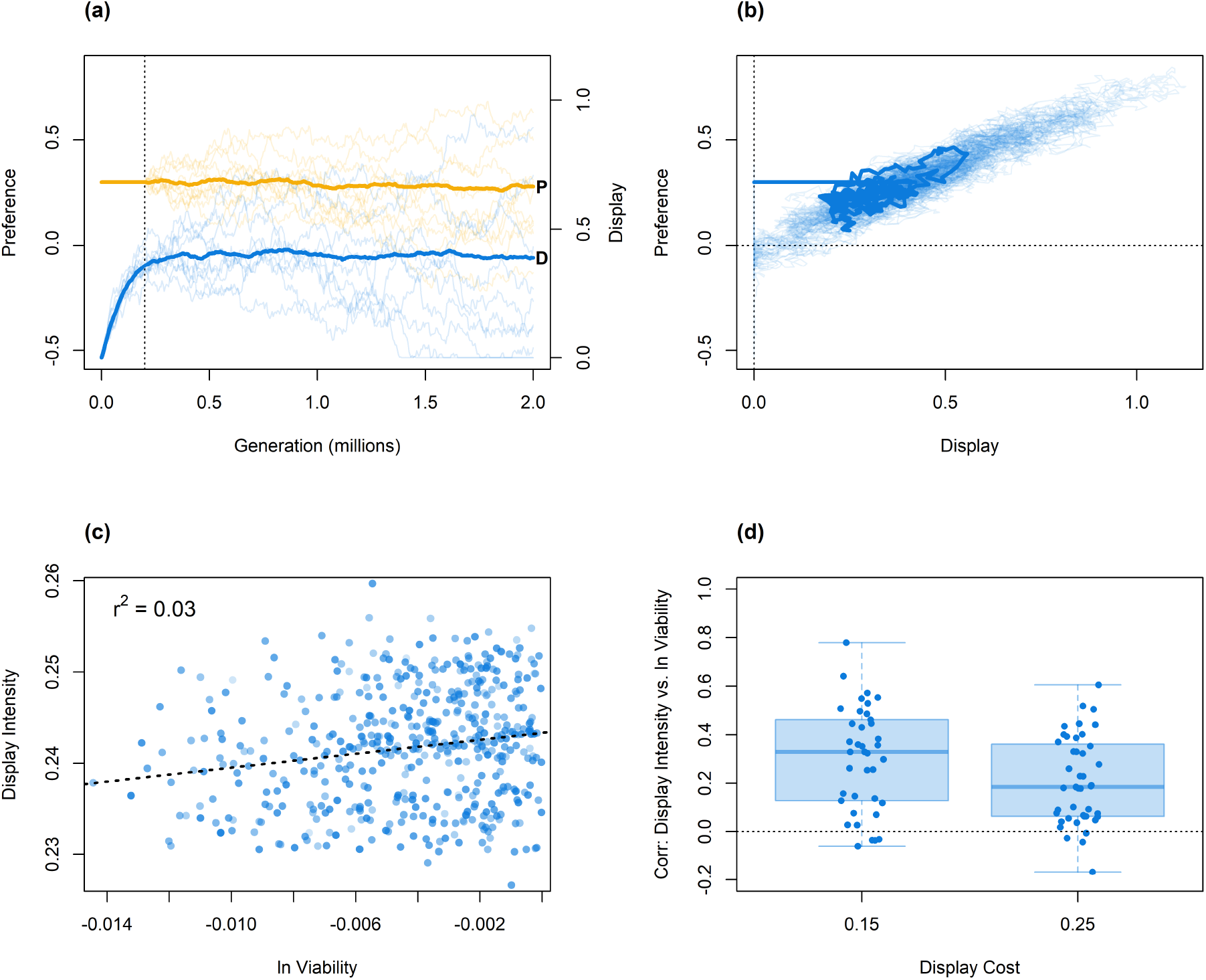
The static display and preference coevolve to positive, stable values. **(a)** Evolution of the static display (blue, **D**) and preference for that display (yellow, **P**). The thin lines show the evolutionary trajectories for 50 individual replicates while the thick lines show the mean value across all replicates. The dotted line marks generation 200,000, past which the preference was allowed to mutate. **(b)** Static display *versus* the preference for that display. The thick line shows the trajectory for the replicate with the median final display value while the light lines show the trajectories for all other replicates. The horizontal and vertical dotted lines mark zero display intensity and zero preference, respectively. **(c)** Display intensity *versus* natural log genetic viability (ln *v* = −|*q*|) for displaying males in the final generation for the replicate with the median final correlation between those two traits. Each data point corresponds to a single display event and shows the associated display intensity and male quality. The dotted line shows the fitted regression line for the two variables. **(d)** Distribution of correlation coefficients (Pearson’s *r*) for display intensity *versus* natural log genetic viability for displaying males in the final generation, for two different display costs. Each data point shows the correlation coefficient for one replicate. The horizontal dotted line marks *r*^2^ = 0. The 19 replicates in which investment was 0 for all display events are not shown, as the correlation coefficient is undefined. Static display cost *β_s_* = 0.25 in (a), (b), and (c), and all other parameter values are as in Table 1.

For the static display to function as an honest indicator of male quality, there must be a positive relationship between displaying males’ genetic viability and realized display intensity. At generation 2,000,000 we observed (Figure 3c,d) a weakly positive relationship (*r*^2^ = 0.04, 95% CI [0.02,0.07], across replicates with *β_s_* = 0.25). We can conclude that static display intensity evolves to be an honest indicator of quality, supporting the evolution of a costly preference for the display, in line with the findings of established models of sexual selection.

### 4.2 Non-flexible, Dynamic Display

The second set of simulations included only the dynamic display (so we set *t_s_* = 0). We further assumed that males’ dynamic display effort did not vary between encounters with females (*d*_2_ = 0). We refer to this display as the *non-flexible, dynamic display*. For these simulations, we set the initial values for the preference and display to *p_d_* = 0.3 and *t_d_* = 0, respectively, for all individuals in the first parental generation. We set the dynamic display cost *β_d_* to two values, 0.015 and 0.025, and ran 50 replicates for each value. We set the values of all other parameters as in Table 1.

As with the static display, the non-flexible, dynamic display and the preference for it evolved to stable, positive values (Figure 4a), with a positive relationship between the two traits (Figure 4b).

**Fig. 4:**
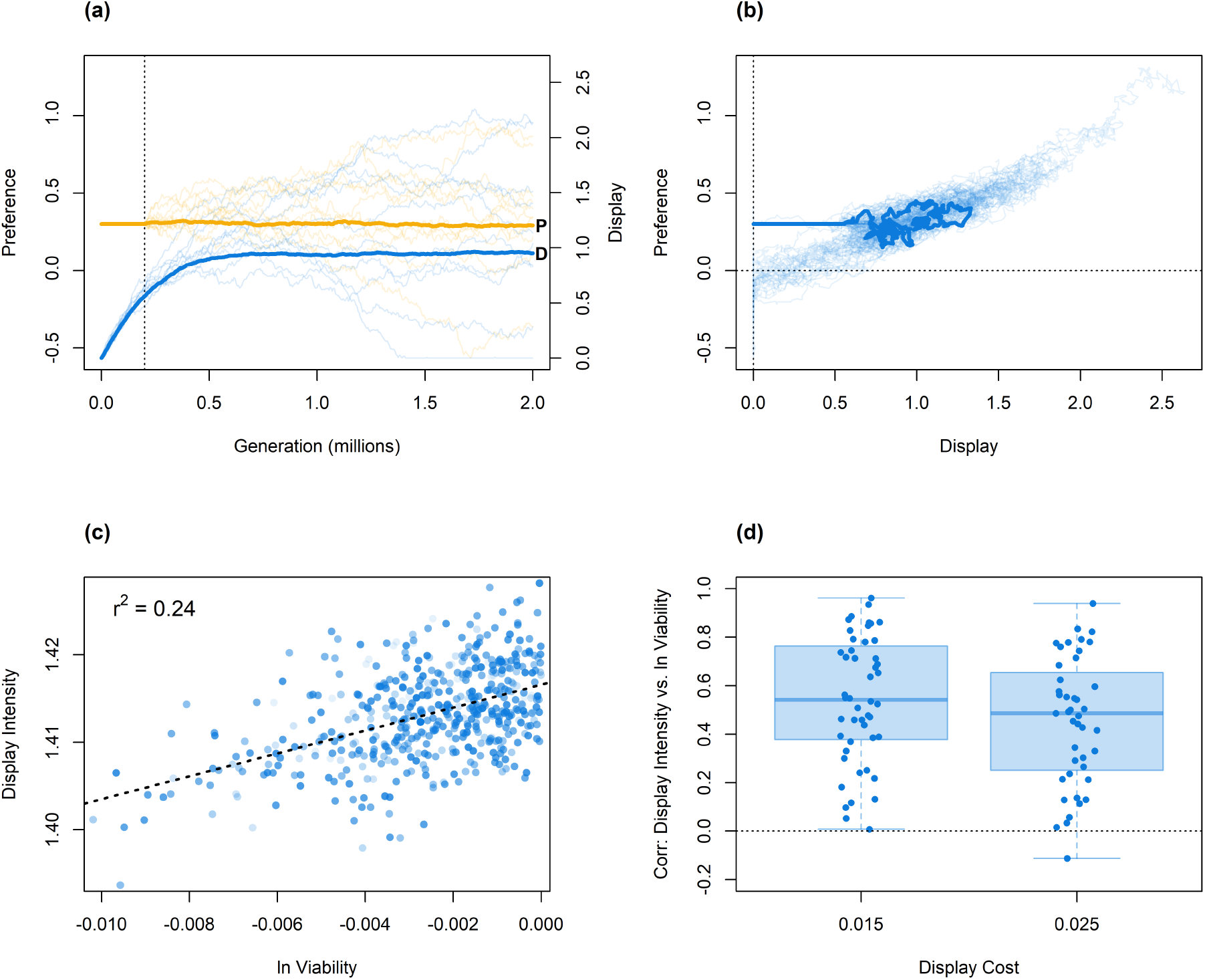
The non-flexible, dynamic display and preference coevolve to positive, stable values, similar to the static display. All data are analogous to those in Figure 3, but for the non-flexible, dynamic display instead of the static display. In **d**, the 9 replicates in which investment was 0 for all display events are not shown, as the correlation coefficient is undefined. Dynamic display cost *β_d_* = 0.025 in (a), (b), and (c), and all other parameter values are as in Table 1.

Note that it is not informative to directly compare the equilibrium values for static and non-flexible, dynamic display intensities. This is because those values depend on the costs of the respective displays, which differ in two key respects. First, on average, males display multiple times and so pay the cost of a dynamic display more than once (hence, we have set *β_d_* an order of magnitude below *β_s_*). Second, males must display at least once before incurring any cost of the dynamic display. Thus, for males that display only once, the dynamic display is no more costly when *β_d_* is low than when *β_d_* is high. For these two reasons, we should not expect *a priori* the values for the static and dynamic displays to evolve to similar levels when the parameters *β_s_* and *β_d_* are similar.

As with the fixed display, we observed a positive relationship (*r*^2^ = 0.21, 95% CI [0.14,0.29], across replicates for *β_d_* = 0.025) between displaying males’ genetic viability and their display intensity at generation 2,000,000 (Figure 4c,d), showing that non-flexible, dynamic display intensity can evolve to be an honest indicator of quality.

These findings are novel but not altogether surprising. They are novel in that no previous model of which we are aware has demonstrated the coevolution of a non-flexible, dynamic display with a sexual preference. Our results show that a non-flexible, dynamic display can evolve to an exaggerated level under a revealing handicap mechanism. The findings are unsurprising because, just as with a static display, each male has a single level of display intensity. Thus, there is no way for lower-quality males to undermine signal honesty by their displaying more intensely under favorable conditions. As such, we should expect a revealing handicap to function with a non-flexible, dynamic display similarly to how it does with a static display: males of lower-quality will consistently display less intensely than those of higher quality.

### 4.3 Flexible Display

The third set of simulations also included only the dynamic display, but we allowed males to flexibly vary their display effort between encounters based on the number of rival males *n* (i.e., *d*_2_ was unconstrained). For these simulations, we again set the initial values for the preference and display to *p_d_* = 0.3 and *t_d_* = 0, respectively, for all individuals in the first parental generation. We set the dynamic display cost *β_d_* to two values, 0.015 and 0.025, and ran 50 replicates for each value. We set the values of all other parameters as in Table 1.

As with the static and non-flexible, dynamic displays, the flexible display and preference for it coevolved to stable, positive values (Figure 5a), with a positive relationship between the two traits (Figure 5b). At the end of the simulations, we observed (Figure 5c,d) a very weakly positive correlation between displaying males’ genetic quality and display intensity (*r*^2^ = 3.3e-4, 95% CI [5.9e-5, 8.1e-4], across replicates for *β_d_* = 0.025). It follows that flexible display intensity evolved to be an honest indicator of quality, but the extremely low *r*^2^ value shows that it is a much noisier indicator than the static and non-flexible, dynamic displays. The likely source of this noise is the evolved strategy for adjusting display investment in response to random variation in the number of competitors.

**Fig. 5:**
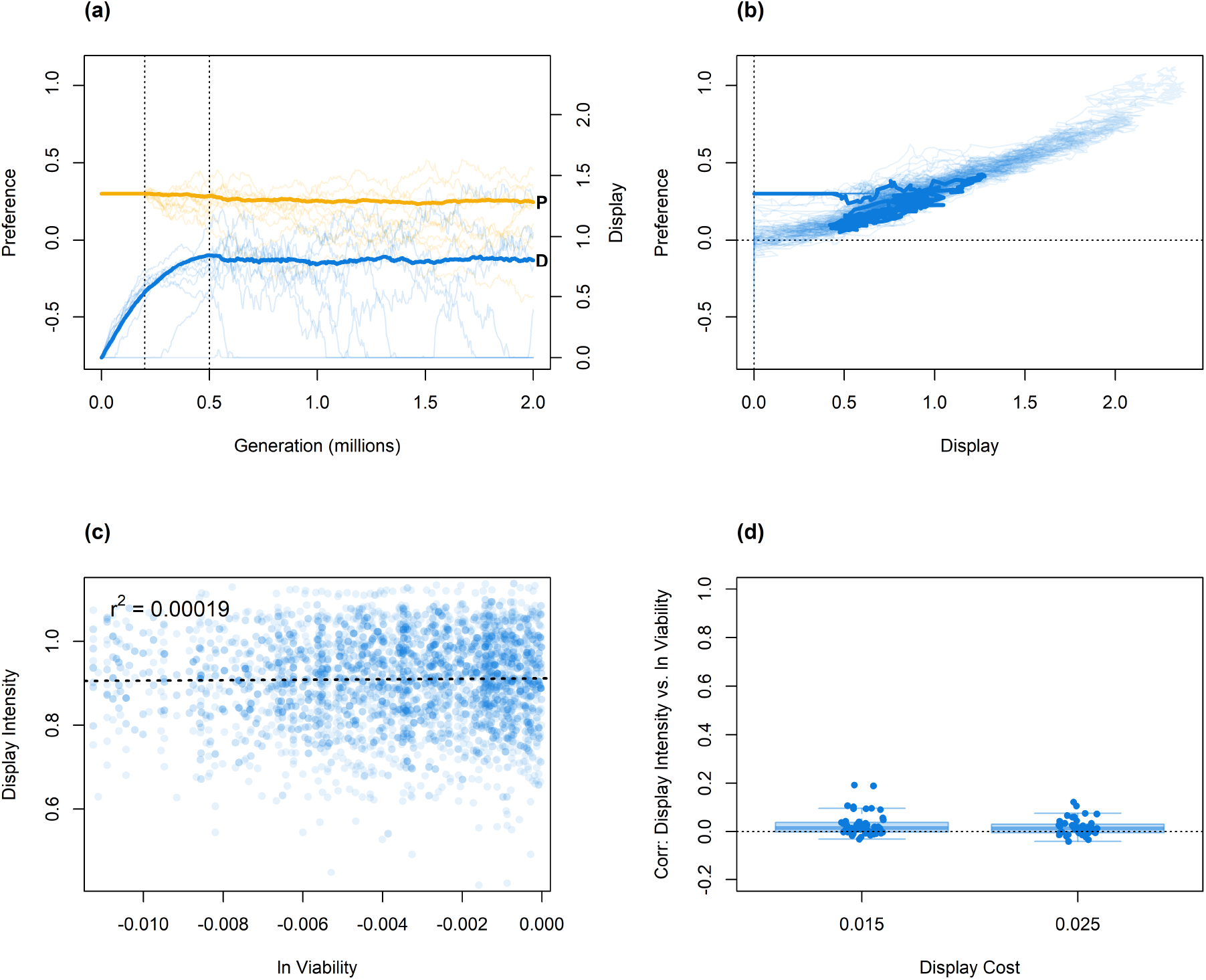
The flexible, dynamic display and preference coevolve to positive, stable values, similar to the static and non-flexible, dynamic displays, but the display is a weaker signal of male quality. All data are analogous to those in Figure 3, but for the flexible display instead of the static display. In **(a)**, the blue lines show males’ investment in the flexible display when displaying alongside the average number of rival males, 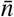; the leftmost dotted line marks generation 200,000, past which the preference was allowed to mutate while the rightmost dotted line marks generation 500,000, past which the display slope was allowed to mutate. In **d**, the 7 replicates in which investment was 0 for all display events are not shown, as the correlation coefficient is undefined. Dynamic display cost *β_d_* = 0.025 in (a), (b), and (c), and all other parameter values are as in Table 1.

The key way that display flexibility might undermine signal honesty is by allowing lower-quality males to invest more heavily in particular signaling contexts so that, if the strength of the female preference is insensitive to the signaling context, as the present model implicitly assumes, then lower-quality males could potentially ”cheat” by displaying more intensely than higher-quality males in those contexts. However, this route to signal dishonesty would require that sufficient genetic covariance accumulate between quality and the genetically determined display investment strategy. In contrast, in the present model, the relationship between quality and realized display intensity approximated the fixed relationship dictated by the revealing handicap mechanism, namely that the display intensity increases exponentially in *v*, as shown in Figure 5c. We further find no relationship between the quality and the dynamic display slope *d*_2_ of displaying males, indicating that there is no systematic difference in display flexibility between low-viability and high-viability males. We can conclude that (a) flexibility does not inherently favor males of lower (or higher) quality and/or (b) whatever advantage flexibility might provide is obscured by recombination preventing the buildup of genetic covariance between quality and display investment.

These results answer the first of our initial questions: a flexible display, where display investment varies with the number of rival males, can evolve under a revealing handicap. Turning to our second question, we now examine the patterns of display flexibility that resulted from the coevolutionary process. Immediately after it is allowed to mutate, the display slope (*d*_2_) falls to negative values and typically continues to decline through to the end of the simulation (Figure 6a). At the end of the simulation, the mean slope is negative with a mean of −0.088, 95% CI [−0.095, −0.080]. Figure 6b depicts the final display function for each replicate. These results suggest that, in the scenario we have modelled, males should reduce their display intensity when displaying alongside more rivals.

**Fig. 6:**
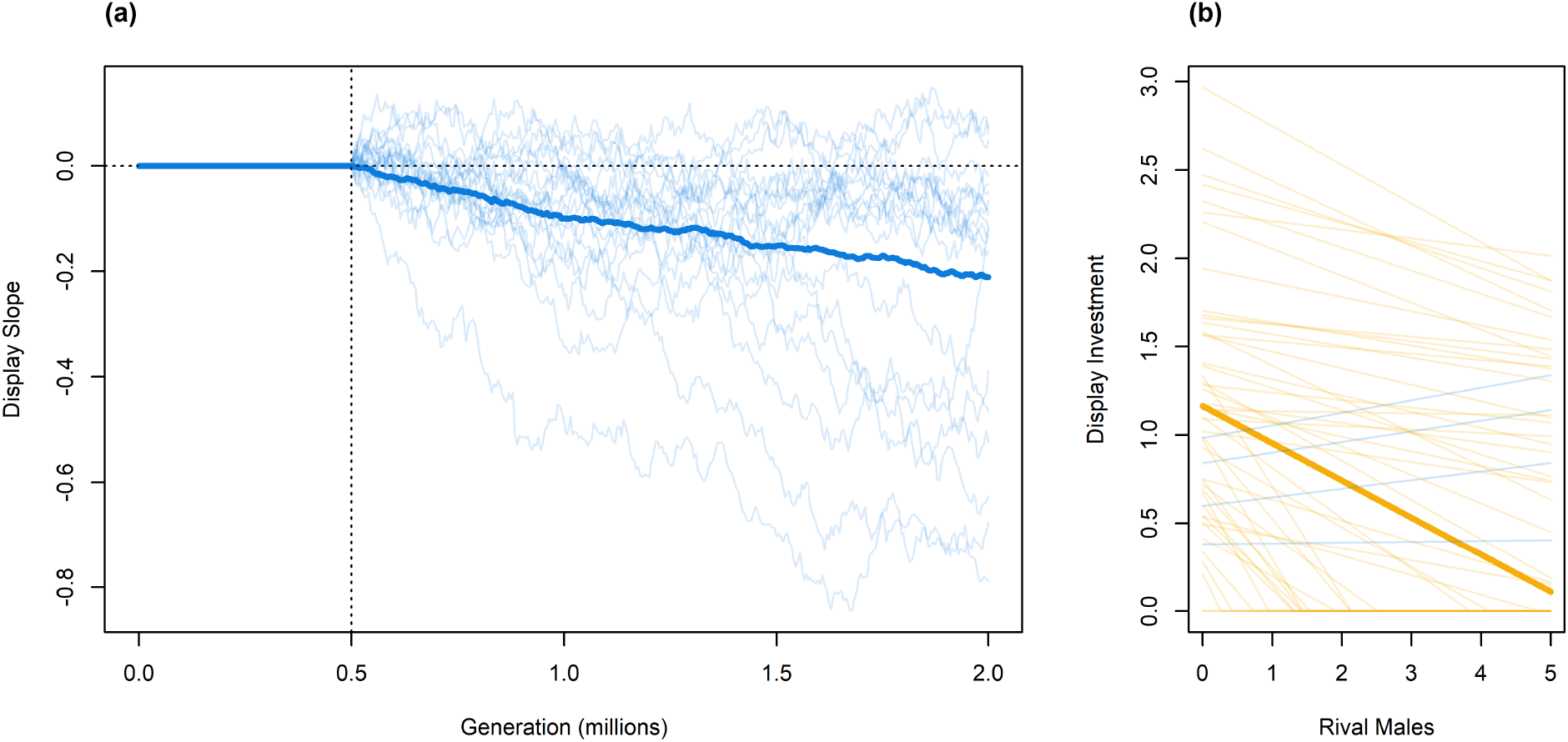
Males evolve to display less intensely when they face more rivals. **(a)** Dynamic display slope *d*_2_ over time. The vertical dotted line marks generation 500,000, past which the display slope was allowed to mutate. The horizontal dotted line corresponds to display investment that is constant with respect to the number of rival males (*d*_2_ = 0). The thick line shows the trajectory for the replicate with the median final display value while the light lines show the trajectories for all other replicates. **(b)** Dynamic display investment *t_d_*(*n*) *versus* number of rival males *n* at the end of the simulation. Each thin line shows the display function for one replicate where the intercept *d*_1_ and slope *d*_2_ are the mean for all males. The thick line shows the display function where *d*_1_ and *d*_2_ are the mean across all replicates. Blue lines have positive slopes (*d*_2_ > 0); yellow lines have negative slopes (*d*_2_ < 0). Note that display functions are horizontal at zero display investment, covering one another. Dynamic display cost *β_d_* = 0.025 in (a), (b), and (c), and all other parameter values are as in Table 1.

## 5 Conclusion

The model presented here is the first to investigate how a sexual preference and a flexible, dynamic courtship display coevolve. We began with an established model of sexual selection (the static-display model) and reproduced the established result that a display trait and sexual preference can coevolve to exaggerated levels under a revealing handicap mechanism. But this version of the model, and the prior models that inspired it, only closely resemble a narrow class of empirically observed courtship displays: those whose expression (and costs) are fixed across an individuals’ lifetime or stages of growth. In reality, even fixed, morphological ornaments (e.g., plumage) are often just one component of more complex, dynamic displays (e.g., a dance) that may alter the ornaments’ perceived intensity. To account for this much broader class of displays whose expression varies between courtship events, we extended the established model to include dynamic and flexible display traits. Our results showed that dynamic displays can coevolve with a sexual preference to exaggerated levels through the same handicap mechanism as a static display. When males’ were allowed to flexibly adjust their display investment based on the number of rivals courting the same female, the display became a far noisier indicator of male quality, but the costly female preference still evolved and still supported costly display expression. The extended model also revealed how males evolved to tactically adjust their display investment: in the scenario we modelled, males decreased their display investment in response to an increased number of rival signalers.

The central significance of this work is that it shows that some key results and insights of established models of static displays generalize to a much larger class of courtship displays. These results, however, should be taken as preliminary—a first step towards generalizing sexual selection theory to account for dynamism and flexibility in display expression. Numerous extensions to the present model that could enhance its biological realism immediately suggest themselves. We restrict ourselves to considering seven.

First, the linear function is just one simple functional form for the flexible display. A key advantage of the linear display function is that it is straightforward to analyze, both because it takes only two parameters and because linearity ensures that a mutation of any given magnitude alters the overall display investment by a fixed amount. (If, for example the display function were instead accelerating in any of its parameters, then the effect of mutations would be greater when the existing investment in the display was high.) The trade-off for this simplicity is reduced flexibility. In the present model, males’ investment can neither accelerate nor decelerate in the number of rivals. To determine the model’s robustness to this restriction, future research could consider adding a curvature parameter *d*_3_ to the display function (e.g., *d*_1_ + *d*_2_*n*^*d*_3_^ rather than *d*_1_ + *d*_2_*n*). Moreover, in the current model, display investment is defined continuously over all numbers of rivals. One effect of this restriction is that males must adjust their investment the same amount between encounters with one rival and two rivals as between encounters with one rival and no rivals. But it seems plausible that males might rather adopt distinct display strategies when displaying alone versus with one or more rivals. Future research could investigate this possibility by adopting a piece-wise display function, with separate parameters determining investment in the two contexts. Ultimately, empirical evidence is required to determine a realistic functional form. For example, male fiddler crabs appear to adjust their waving rate little between one and two waving neighbors and then increase their waving rate approximately linearly with higher numbers of rivals (Milner et al., 2012).

Second, we have assumed that the same genes determine display expression in males of differing qualities. If instead the genes that determine quality also determined which genes coding for display were expressed, then it would be possible for low-quality display investment and high-quality display investment to evolve as separate traits. This would allow signaling strategies that deviate more from the fixed relationship dictated by the revealing handicap. For example, low-quality display investment could evolve to a higher level than high-quality display expression, partially or fully counteracting the effects of the revealing handicap. Modeling this scenario would allow a more thorough investigation of the conditions under which the revealing handicap supports signal honesty (van Doorn & Weissing, 2006) for static, dynamic, and flexible displays.

Third, we have assumed that females do not adjust preference strength by the number of rival males. Yet if males can perceive the number of rivals they display alongside and respond accordingly, it seems plausible to assume that females could use the same information. If females adjusted the strength of their preferences to counteract males’ increased (or reduced) display effort, this could potentially reduce the fitness advantage of displaying flexibly.

Fourth, we have modeled the display as either purely static or purely dynamic, but most displays in nature have both static and dynamic components (Patricelli et al., 2016). Past models suggest that it is difficult to sustain costly preferences for multiple indicators of the same aspect of quality (van Doorn & Weissing, 2004), so given the dynamic display’s poorer information content, we might expect the dynamic display to be lost and the static display retained if the two were initially active in the present model as separate, non-interacting traits. However, the two displays could plausibly synergize with one another as part of a more complex, overall display, such as when a dance shows off bright plumage. The model could be extended to account for this scenario by including both the static and the dynamic display, *t_d_* and *t_s_*; the female preferences for both displays, *p_s_* and *p_d_*; and a preference *p_x_* for the interaction between the static and dynamic components (i.e., *t_s_ × t_d_*). Then, the display could evolve to be completely static (*t_d_* = 0), completely dynamic (*t_s_* = 0), or in between (*t_s_* ≠ 0, *t_d_* ≠ 0), with females potentially preferring a level of dynamism anywhere along that spectrum.

Fifth, we have assumed that the flexible display imposes a cost only through decreased viability. However, when multiple displays take place within a short time frame, investing in the display might also reduce the signaler’s condition, impairing further displays for a limited time. Changes in current condition would then introduce extra variation in the display intensity of a male of any given quality. Then we might expect condition-dependent signaling strategies to evolve, such that males might compensate for their fatigue by increasing their investment or conserve their energy by decreasing their investment. A state-dependent model would be needed to analyze such strategies and would differ fundamentally from the viability-based model developed here, in that the fatigue induced by the dynamic display (whether non-flexible or flexible) could wear off over time, and hence the cost of the display would depend crucially on investment in recent display events.

Sixth, the present model does not incorporate signal interference or male-male competition, both of which may affect the evolution of flexible display strategies. For example, male bird-voiced tree frogs, *Hyla avivoca*, modify their calls to avoid overlap with the calls of rivals (Martínez-Rivera & Gerhardt, 2008), indicating that acoustic interference may be an important driver of display flexibility. As another example, male sage grouse have been observed to avoid signaling when doing so would risk a fight with rival males present at a lek (Patricelli et al., 2016), suggesting contests and injury as another potential driver of display flexibility.

Seventh, we have allowed males to adjust their display investment based only on the number of rivals displaying. But males might plausibly adjust their displays based on a variety of other cues. For example, when females vary in quality or fecundity, males might alter their investment based on cues to those traits, increasing their courtship effort for more desirable mates. Alternatively, if displaying attracts predators, males might plausibly display more intensely when and where the risk of predation appears lower. Another possibility is that, when the population density varies over time, males might adjust their display intensity based on the frequency of encounters with potential mates. Yet another possibility is that males might display more intensely when alternative activities, such as food gathering, are less worthwhile, such as in poor light conditions. For example, such variations in the opportunity cost of displaying may explain why birdsong often peaks at twilight (Hutchinson et al., 1993). To our knowledge, no existing model of the coevolution of a display and preference has incorporated any of these sorts of tactical display adjustment.

We have, in sum, focused entirely on the display flexibility that arises due to female choice between varying numbers of rival males and showed, for the first time, that trait exaggeration can occur with such flexibility. The vast array of other empirically documented and theoretically compelling sources of within-individual display variation outlined above—yet to be integrated into established sexual selection theory—await modeling. After decades of theory have revealed a rich picture of the evolution of static sexual ornaments, it is time to generalize the field’s findings to account for the ubiquitous dynamism of animal courtship.

## 6 Statements and Declarations

### 6.1 Author Contributions

TWF and IGR conceived the study and built an initial version of the model. SH developed the full model with input from TWF and TV, analyzed the output, and wrote the first draft of the manuscript. All authors commented on the manuscript and read and approved the final version.

